# Methionine uptake via SLC43A2 transporter is essential for regulatory T lymphocyte survival

**DOI:** 10.1101/2022.03.09.483598

**Authors:** Afsana Naaz, Neetu Saini, Shree Padma, Pinki Gahlot, Adhish Walvekar, Anupam Dutta, Umamaheshwari Davathamizhan, Apurva Sarin, Sunil Laxman

## Abstract

It is increasingly clear that cell death, survival or growth decisions of T lymphocyte subsets depend on interplay between cytokine-dependent and metabolic processes. What the metabolic requirements of T regulatory cells (Tregs) for their survival are, and how these requirements are satisfied remain to be fully understood. In this study, we identified a necessary requirement of methionine uptake and utilization for Tregs survival upon interleukin 2 (IL-2) deprivation. Activated Tregs have high methionine uptake and consumption to S-adenosyl methionine (SAM) and S-adenosyl homocysteine (SAH). This methionine uptake is essential for Tregs survival, and is regulated by Notch1 activity. Notch1 controls the expression of the solute carrier protein SLC43A2 transporter during IL-2 deprivation. SLC43A2 is necessary for sufficient methionine uptake, and determines Tregs viability upon IL-2 withdrawal. Collectively, we identify a specifically regulated mechanism of methionine import in Tregs that is necessary for the survival of these cells. This highlights the need for methionine availability and metabolism in contextually regulating cell death in an immunosuppressive population of T lymphocytes.

## Introduction

T lymphocytes play central roles in adaptive immune responses. In order to appropriately develop adaptive immune responses, the many T cell sub-classes adapt to a range of extracellular nutrient levels and environmental cues. Several signals push T cells out of quiescence, and towards acquiring new functions. Here, it is clear that metabolic reprogramming is central to the survival, differentiation or function of T cells, working in concert with key signaling systems and cytokine function (1–5). How these are different in distinct T cell sub-types, and how these metabolic requirements are met remain unclear.

Depending upon the T cell subset, distinct nutrients such as glucose, amino acids, and lipids control metabolism, which influences immune signaling and regulates the function and proliferation of T cells (1, 2, 6). In general, T cell activation in response to an antigen results in transcriptional and metabolic remodeling, leading to new functions including the production of cytokines and molecules that support T cell survival, expansion, and functional demands (1, 7–11). Since T cell subsets function in diverse and dynamic niches they require different energetic and metabolic pathways for their survival and function (12–15). For example, upon antigen activation CD4^+^ T effector cells switch from oxidative phosphorylation (OXPHOS) to glycolysis to meet high metabolic demands, while antigen activated CD4^+^CD25^hi^FoxP3^+^ T regulatory cells (Tregs) switch to high lipid oxidation and low glycolysis (3, 15, 16). The differing use of biosynthetic pathways distinguishes T-cell fate choices. In this context, what are the metabolic requirements that control Tregs survival, and how they are met and regulated remains an interesting yet poorly explored area.

Within this framework, distinct amino acids have unique roles in controlling T cell function. Amino acids have diverse roles in metabolism and signaling, and control multiple cellular programs. Studies in naïve T cells noted substantial changes in amino acid pools compared to activated T cells (17, 18). Several amino acids such as leucine, glutamine, arginine, and tryptophan have been identified to regulate T cell homeostasis and function, and the expression of distinct amino acid transporters is critical for this function (19–23). In T effectors (Teffs), critical roles for neutral and branched-chain amino acids were identified in T cell expansion, mTORC1 activation and meeting bioenergetic requirements (23–25). In Teffs, the solute carrier transporter SLC7A5 (which transports large neutral amino acids) was specifically upregulated and required for their expansion in amino acid dependent contexts (24, 26). CD4^+^ and CD8^+^ T cells deficient in SLC7A5 showed impaired clonal expansion and effector functions (24, 26), and reduced mTORC1 activation concurrent with decreased glutamine and glucose uptake following T cell activation (24). More recently, multi-faceted roles for methionine were identified for T cell activation and differentiation/expansion into Teffs (25). In this context, methionine-transport regulated by the SLC7A5 transporter was the rate-limiting, essential factor in the proliferation and differentiation of these T cells (25). Similarly, branched-chain amino acids transported by SLC3A2 and CD98 regulates activation and suppressor function in Tregs (27, 28). These data suggest the likely existence of many as yet unknown roles of amino acids and amino acid transporters in regulating T cell fates.

Tregs are a sub-class of T cells that are crucial for peripheral tolerance and immune homeostasis (29, 30). Their immunosuppressive function is critical for pathologies such as autoimmunity, cancer and tissue damage (31, 32). Tregs have unique metabolic requirements that are distinct from other T cells like the Teffs (3, 33, 34). Tregs must survive and protect themselves in dynamic and nutrient limiting microenvironments, and these survival decisions have been best studied in the context of cytokine requirements in these cells. Therefore, understanding how Tregs regulate survival/death programs becomes critical in order to address how they function. Relevantly, activated CD4^+^ and CD8^+^ T effectors require cytokines like IL-2 for their survival, and undergo apoptosis upon its withdrawal (35, 36). However, Tregs continue to survive in cultures without IL-2 for extended periods of time (37), and this indicates that additional, as yet unknown, factors enable the survival of Tregs. Here, non-canonical (cytoplasmic) Notch1 activity controlled Tregs survival, by regulating mitochondrial activity, and metabolism (37, 38). Unlike in Teffs where Notch1 is localized in the nucleus, in Tregs Notch1 is predominantly cytoplasmic, and carries out its protective effects (37). While these roles of Notch1 in enabling Tregs survival are critical, the unique metabolic requirements of Tregs for their survival, and how these metabolic requirements are satisfied, remain unknown.

Herein, we report unique amino acid requirements for Tregs survival when IL-2 is deprived. Upon IL-2 withdrawal, Tregs cells exhibit an increased requirement of a single amino acid, methionine, which is critical for cell survival. This is enabled by the Notch1 mediated regulation of the activity of a specific solute carrier transporter. The transporter mediated uptake and thereby utilization of methionine is the limiting factor for Tregs survival. We thus identify an essential amino acid requirement for Tregs survival upon IL-2 withdrawal. This is mediated by a novel Notch1-regulated amino acid transporter axis that regulates the sustained supply of methionine and thereby Tregs survival.

## Results

### Activated Tregs uptake and metabolize methionine upon cytokine withdrawal

To investigate the status of amino acids in Tregs survival contexts, we measured steady state amino acid pools in primary murine Tregs upon IL-2 withdrawal. Intracellular amino acids were assessed over the first ~6 h after IL-2 withdrawal in Tregs, while they remained in complete medium with dialyzed serum (CMDS). We used targeted, quantitative LC/MS/MS based approaches to investigate the intracellular amino acid abundance in these cells. Metabolites were extracted from activated Tregs cultured in CMDS without IL-2 for 1, 3, and 6 h (Fig 1A: inset schematic), and intracellular levels of amino acids were assessed. A majority of the amino acids did not show significant changes in their amounts (Fig 1A) (indicating an overall stable homeostasis with the external environment). Contrastingly, intracellular methionine level showed the most substantial decrease, as rapidly as within 1 h of cytokine withdrawal, along with a decrease in the related sulfur-amino acid cysteine, as well as histidine and tryptophan (Fig 1A). This suggested a shift in homeostasis in these cells, towards increased methionine uptake and consumption. Methionine is converted into its major metabolite SAM, and methyltransferases transfer the methyl group from SAM to produce a methylated substrate and SAH. We therefore assessed the relative amounts of the products of methionine utilization, SAM and SAH, as part of the methionine cycle (Fig 1B). IL-2 withdrawal altered SAM pools (Fig 1B), and thus the SAM/SAH ratios in Tregs, consistent with increased SAM consumption. These data suggested a possible acute requirement of methionine in Tregs, upon IL-2 withdrawal. To unambiguously determine whether this requirement was provided by extracellular methionine that is taken up by Tregs, we studied the uptake and utilization of 13C5N15 labeled methionine provided in the cell culture medium. This was done by collecting metabolites from Tregs cultured in sulfur amino acid (SAA) dropout medium (DOSAA) and without IL-2 for 1, 3, and 6 h. Labeled methionine was added to cells 20 min before metabolite extraction (Fig 1C). Tregs continuously took up and utilized methionine (post IL-2 withdrawal), as indicated by the ~100% label incorporation (and no remaining unlabeled methionine coming from existing stores in the cell) even after this brief time-window of label addition (Fig 1D (i)). Further, the amount of label incorporation into SAM and thereby SAH, which indicates the extent of methionine consumption, was highest 1 h after IL-2 withdrawal, and decreased over 3 h (Fig 1D (ii)).

**Figure 1.**
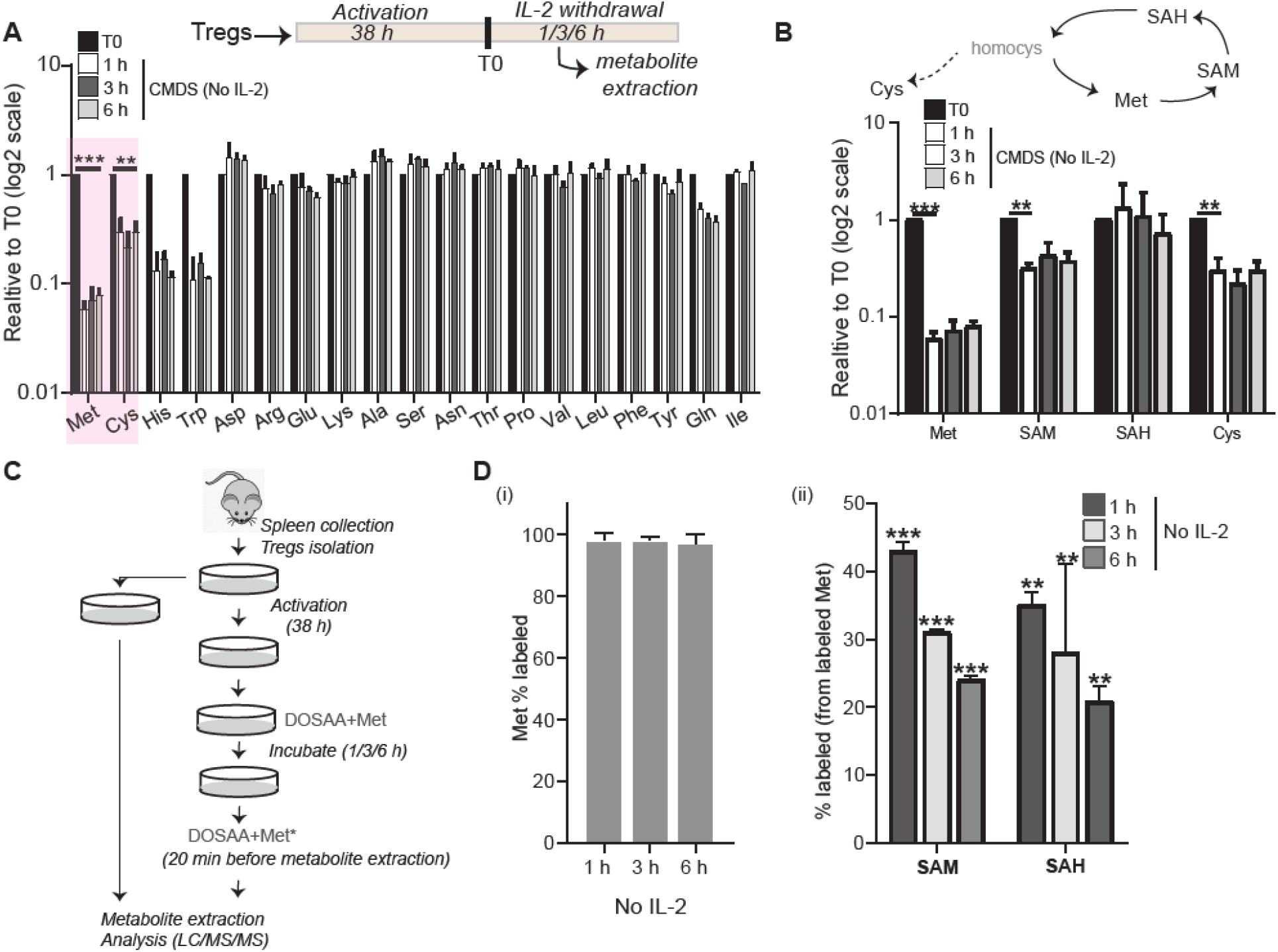
Tregs have a specific uptake and utilization of methionine upon IL-2 withdrawal. (A) Mass spectrometric analysis to determine relative changes in intracellular amino acid amounts in Tregs cultured in CMDS without IL-2 for 1, 3, and 6 h. Amino acid amounts are plotted relative to T0 (onset of IL-2 withdrawal). Inset Schematic: Experimental timeline for assessment of metabolites in activated Tregs. (B) Intracellular levels of methionine, SAM, SAH and cysteine in Tregs cultured in CMDS without IL-2 for 1, 3, and 6 h. (C) Experimental protocol to assess methionine uptake in activated Tregs, using 13C5N15 labeled methionine. Methionine (150 μM) was added at T0 and labeled methionine (150 μM) was added 20 min prior to metabolite extraction for each time point. (D) Targeted LC/MS/MS based analysis to assess the relative uptake and utilization of extracellular (i) 13C15N labeled methionine and subsequently labeled (ii) SAM and SAH in Tregs cultured in DOSAA without IL-2 for 1, 3, and 6 h. Significance value calculated with respect to T0. Data indicates Mean ± SD of two independent experiments. ***p≤0.001 and **p≤0.01.

Collectively, these data demonstrate that Tregs take up extracellular methionine, consume it to synthesize SAM, and utilize SAM for methylation reactions, with the highest uptake and utilization over the first ~1 h after IL-2 withdrawal.

### Methionine is essential for the survival of Tregs in the absence of IL-2

Activated Tregs can survive after IL-2 deprivation (37). To further investigate if methionine had any role in this survival, Tregs were cultured without IL-2 either in SAA dropout or complete media with dialyzed serum. The dropout of SAAs abrogated Tregs survival in the absence of IL-2 (Fig 2A). However, adding methionine at the onset of the assay (T0) rescued cells from apoptosis triggered by IL-2 withdrawal. Cysteine, unlike methionine, did not confer protection from apoptosis to Tregs cultured in the absence of SAAs and IL-2 (Fig 2A). Further, cysteine in combination with methionine did not provide additional protection to Tregs (Fig 2A). These data suggest that the protective role is specific to methionine. To further test this methionine requirement; in complementary experiments we supplemented cells with ethionine, an antimetabolite and antagonist of methionine, which would competitively block methionine uptake and utilization. Ethionine strongly inhibited the methionine-dependent cell survival in IL-2 starvation (Fig 2B). Finally, the protection of Tregs from apoptosis was greatest when methionine was added back at the initial h (between 0-5 h, i.e., T0 and T5) as compared to ~9 h after IL-2 withdrawal (T9) (Fig 2C). These data suggest a specific, early requirement of methionine uptake and utilization for Tregs survival post IL-2 withdrawal.

**Figure 2.**
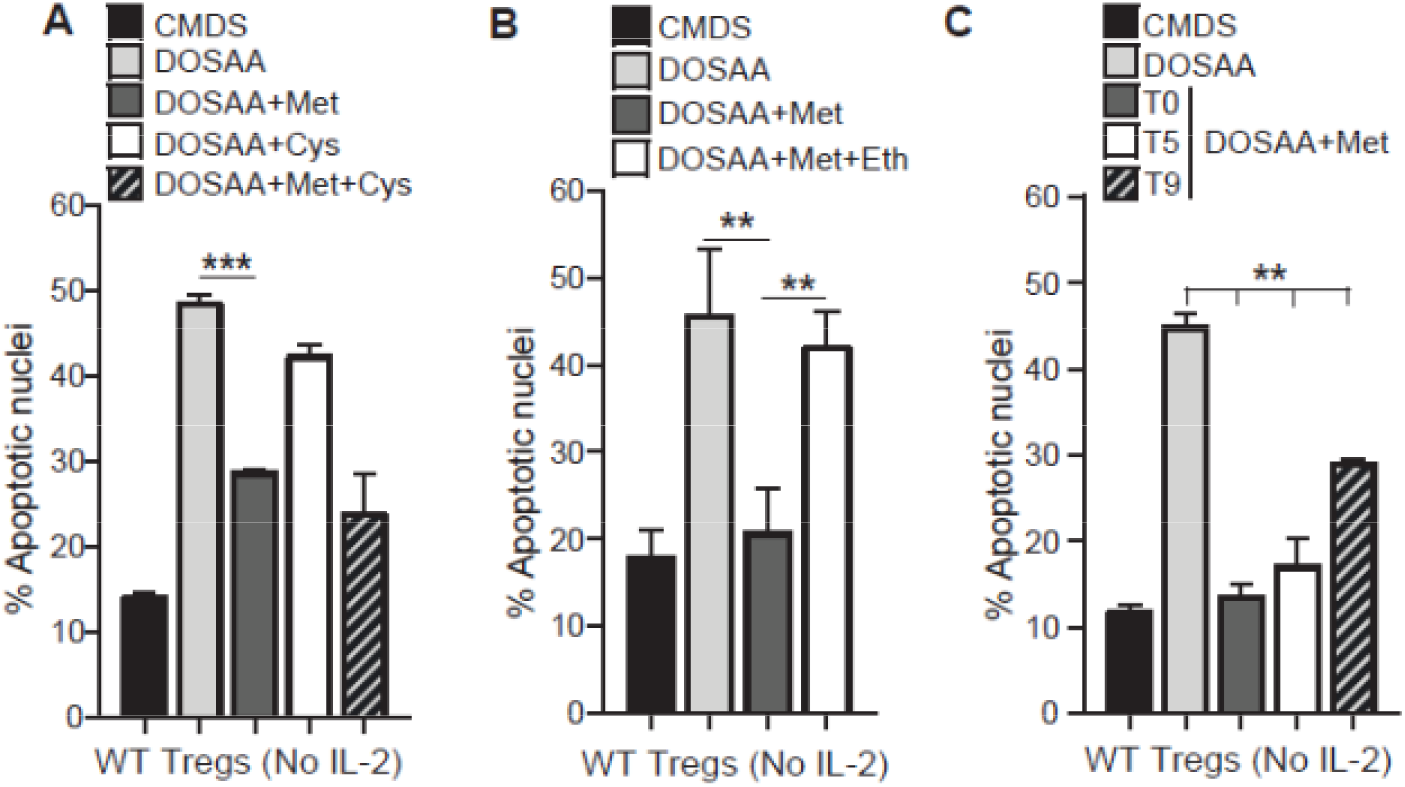
Activated Tregs require methionine for survival upon IL-2 withdrawal. (A) Apoptotic nuclei in WT Tregs cultured for 18-22 h in CMDS and DOSAA without and with methionine (150 μM) and cysteine (150 μM) added separately or in combination. Methionine and cysteine were used at a concentration of 100 μM in combination. (B) Apoptotic nuclei in WT Tregs cultured for 18-22 h in CMDS and DOSAA without and with methionine (150 μM) added at T0, or methionine in combination with ethionine (750 μM). Ethionine was added at T0 followed by addition of methionine after 20 min. (C) Apoptotic nuclei in WT Tregs cultured for 18-22 h in CMDS and DOSAA with methionine (150 μM) added at T0 or after 5 h (T5) and 9 h (T9) of the onset of the assay. Data indicates Mean ± SD of two independent experiments in panels A and C, and three independent experiments in panel B. ***p≤0.001 and **p≤0.01.

### Notch1 function is required for methionine uptake and survival of Tregs upon cytokine withdrawal

In IL-2 limiting conditions, Notch1 (non-nuclear) enables the survival of Tregs (37). We therefore asked if the methionine-dependent survival of Tregs following IL-2 withdrawal also requires Notch1 activity. GSI (GS inhibitor-X), a pharmacological inhibitor of the enzyme γ-secretase was used to inhibit cleavage and release of the Notch1 intracellular domain (37). The inhibition of Notch1 activity by GSI significantly reduced the protection conferred by methionine on Tregs survival (Fig 3A). We next asked if the methionine-dependent protection of Tregs apoptosis is affected in the absence of Notch1. For this, we utilized Tregs isolated from Notch1^+/+^ (Cre-ve) or Notch1^−/−^ (Cre+ve; Cd4-Cre∷Notch1^lox/lox^) mice. The addition of methionine protected activated Notch1^+/+^ (Cre-ve) Tregs from apoptosis when cultured in the absence of SAAs and IL-2 (Fig 3B). However, methionine-addition failed to rescue the survival of activated Tregs obtained from mice with the targeted ablation of Notch1 (Notch1^−/−^) and cultured without IL-2 (Fig 3B). These data show that Notch1 function is required for the methionine-mediated Tregs survival post IL-2 withdrawal. To further understand if Notch1 had any role in mediating methionine uptake in Tregs, we assessed the change in the mRNA levels of several amino acid transporters following IL-2 withdrawal, in the presence and absence of GSI. These transporters belong to the solute carrier (SLC) superfamily, which are the major transporters of several amino acids including methionine (39). Among the transporters tested, SLC6A17, SLC1A5, and SLC7A8 showed very high Ct values (>30) (Fig 3C, Table S1) indicating very low/basal transcript levels in Tregs. In contrast, SLC3A1, SLC43A1, SLC43A2 and SLC7A5 all had higher mRNA expression (Fig 3C). Interestingly, the removal of IL-2 appeared to increase the expression of SLC43A2 and SLC43A1 (Fig 3D). Further, abrogating Notch1 activity in Tregs cultured without IL-2 decreased the mRNA levels of the SLC43A transporters (Fig 3D). None of the other SLC transporters revealed any trend of putative Notch1 dependent expression. Notably, the mRNA levels of SLC7A5, a transporter required for the import of methionine in CD4^+^ T cells (25) remain unaltered in Tregs. These data collectively suggested that the SLC43A transporters might play a role in the Notch1 mediated regulation of methionine uptake in Tregs.

**Figure 3.**
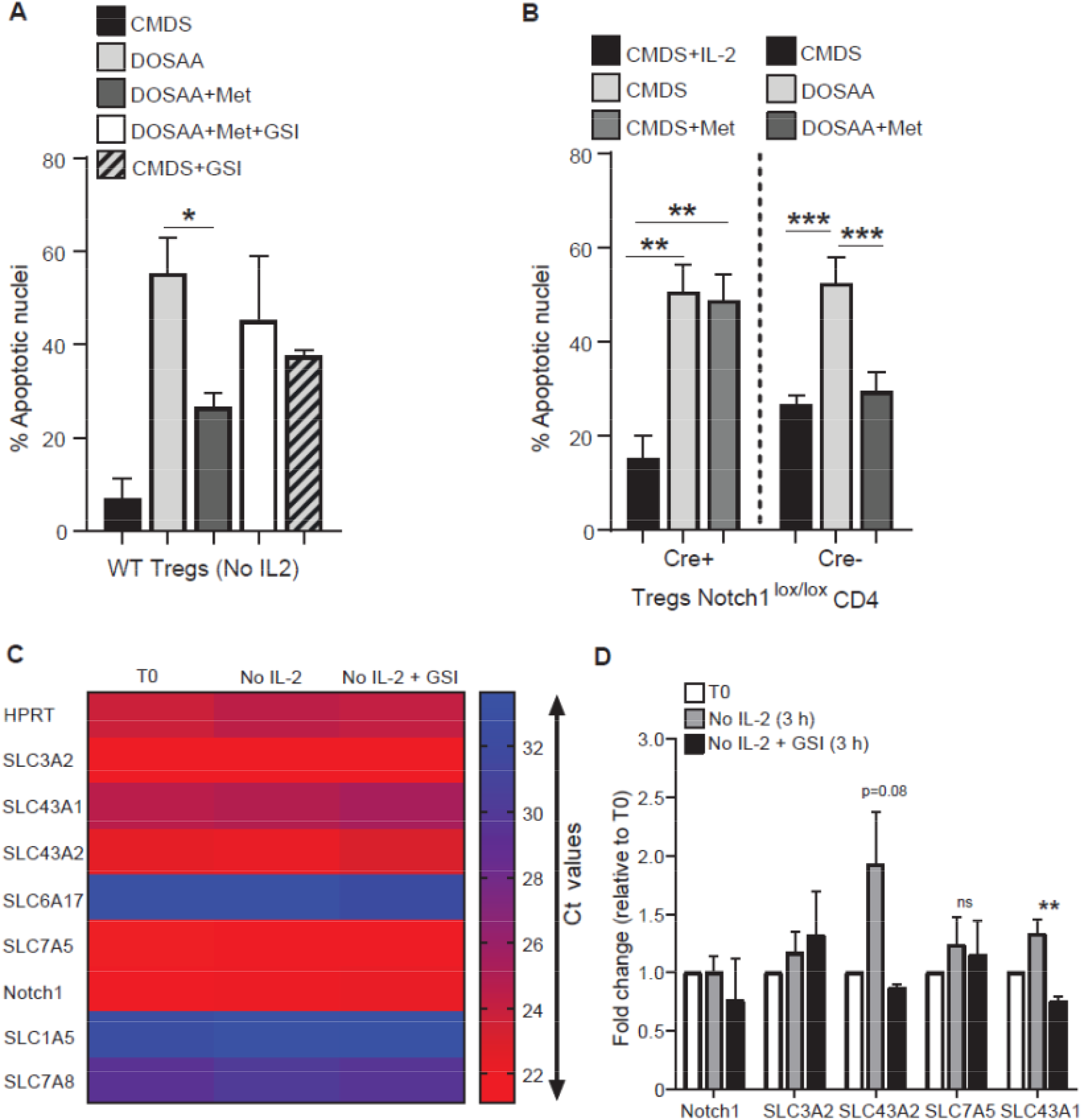
Notch1 regulates methionine-dependent Tregs survival. (A) Apoptotic nuclei in Tregs cultured for 18-22 h in CMDS and DOSAA without or with methionine (200 μM) added at T5 either in the presence (7.5 μM) or absence of GSI. (B) Apoptotic nuclei in Notch1^−/−^ (Cre+ve; Cd4-Cre∷Notch1^lox/lox^)Tregs (left) when cultured for 18-22 h in CMDS, CMDS with IL-2, and CMDS with methionine (150 μM) and apoptotic nuclei in Notch1^+/+^ (Cre-ve) Tregs (right) when cultured for 18-22 h in CMDS and DOSAA without and with methionine (150 μM). (C) Heat map showing Ct values of indicated SLC amino acid transporters in Tregs at T0 and post 3 h culture without IL-2 in the presence (10 μM) or absence of GSI. (D) Fold change in mRNA levels of indicated SLC amino acid transporters in Tregs cultured for 3 h without IL-2 in the presence (10 μM) or absence of GSI. Fold change was measured relative to T0. Data indicates Mean ± SD of two independent experiments for panel A, C and D and three independent experiments for panel B. ***p≤0.001, **p≤0.01 and *p≤0.05, ns implies non-significant.

### SLC43A2 transporter is essential for methionine dependent Tregs survival

The SLC43A2 protein had high transcript amounts, as well as Notch1 dependent expression. We therefore prioritized SLC43A2, and sought to understand if this transporter is required for Tregs survival upon IL-2 withdrawal, in the context of methionine availability. We also assessed if SLC7A5, which regulates the import of methionine in CD4^+^ T cells (25), had any role to play in Tregs survival in the absence of IL-2. Using shRNA, we knocked-down respectively SLC43A2 and SLC7A5 in Tregs (Fig4A(i)). Assessing Tregs survival after IL-2 withdrawal, we found that shRNA to SLC43A2, not SLC7A5 or scramble control abrogated Tregs survival in these conditions (Fig 4A(ii)). This suggests a necessary role for SLC43A2 in Tregs survival upon IL-2 withdrawal. We next asked whether the SLC43A2 transporter protein was required for the methionine-dependent survival of Tregs in IL-2 limiting conditions. Tregs were infected with scramble (control) and SLC43A2 shRNA (Fig 4B(i)). As compared to scramble control, the knock-down of SLC43A2 abrogated Tregs survival when cultured in CMDS, and even when DOSAA was supplemented with methionine (Fig 4B(ii)). These data therefore show that the SLC43A2 transporter has a specific, necessary role in Tregs in enabling the methionine-dependent survival post IL-2 withdrawal.

**Figure 4.**
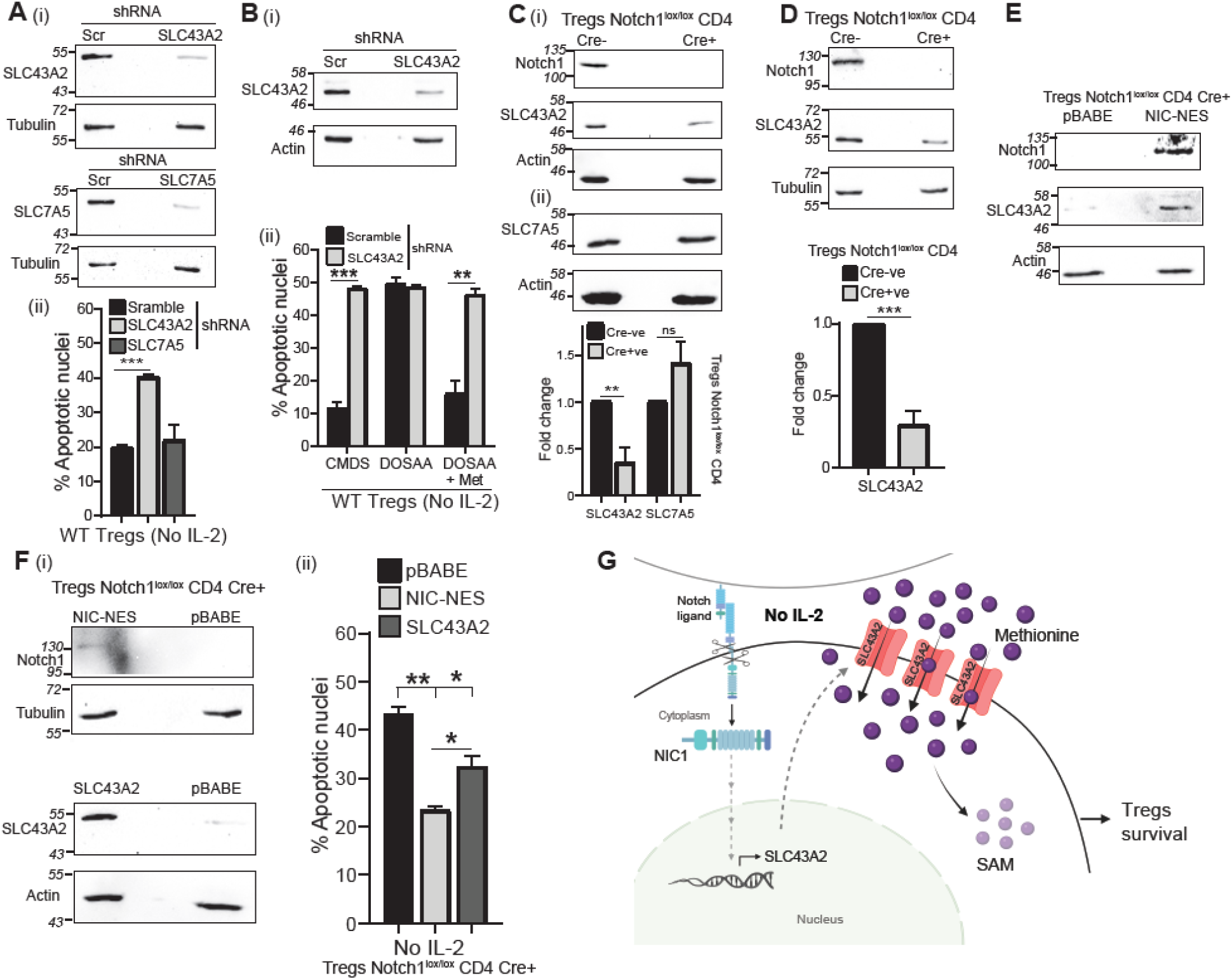
Notch dependent SLC43A2 transporter regulates methionine-dependent Tregs survival. (A-i) Immunoblots probed for SLC43A2 or SLC7A5 and Tubulin in whole cell lysates from WT Tregs transduced with retroviruses containing shRNA to SCL43A2, SLC7A5 or scrambled (Scr) control. Immunoblots are representative of two independent experiments. (A-ii) Apoptotic nuclei in WT Tregs transduced with retroviruses containing shRNA to SCL43A2, SLC7A5 or scrambled control and cultured without IL-2 for 18-22 h. (B-i) Immunoblots probed for SLC43A2 and Actin in whole cell lysates from WT Tregs transduced with retroviruses containing shRNA to SLC43A2 or scrambled (Scr) control. (B-ii) Apoptotic nuclei in WT Tregs transduced with retroviruses containing shRNA to SLC43A2 or scrambled control and cultured in CMDS or DOSAA with (150 μM) or without methionine for 18-22 h. (C) Immunoblots of cell lysates prepared from activated Notch1^−/−^ (Cre+ve; Cd4-Cre∷Notch1^lox/lox^) and Notch1^+/+^ (Cre-ve) Tregs probed for (i) SLC43A2, Notch1 and Actin, and (ii) SLC7A5 and Actin. The cell lysate was split and used for C-i and C-ii. Inset: Densitometric analysis indicating fold change in the protein levels of SLC43A2 and SLC7A5 in Notch1^−/−^ Tregs relative to Notch1^+/+^ Tregs. Data indicates Mean ± SD of three independent experiments for C-i and two independent experiments for C-ii. (D) Immunoblots probed for SLC43A2, Notch1 and Tubulin in freshly isolated Notch1^+/+^ and Notch1^−/−^ Tregs. Inset: Densitometric analysis indicating fold change in the protein levels of SLC43A2 in freshly isolated Notch1^+/+^ and Notch1^−/−^ Tregs. (E) Immunoblots probed for SLC43A2, Notch1 and Actin in Notch1^−/−^ Tregs transduced with retroviruses containing pBABE-NIC-NES or control pBABE plasmid (Immunoblots are representative of three independent experiments). (F-i) Immunoblots probed for Notch1 and Tubulin, and SLC43A2 and Actin in Notch1^−/−^ (Cre+ve; Cd4-Cre∷Notch1^lox/lox^) Tregs transduced with retroviruses containing pBABE-NIC-NES, pBABE-SLC43A2 or control pBABE plasmid. (F-ii) Apoptotic nuclei in Notch1^−/−^ Tregs transduced with retroviruses containing pBABE-NIC-NES, pBABE-SLC43A2 or control pBABE plasmid and cultured without IL-2 for 18-22 h. (G) Model: Upon IL-2 depletion, Tregs continuously take up and utilize methionine for survival. This occurs through the SLC43A2 transporter in a Notch1 dependent manner. Notch1 regulates SLC43A2 expression, which increases upon IL-2 depletion. This allows Tregs to take up methionine, and utilize it, enabling Tregs survival. Data indicates Mean ± SD of two independent experiments in panels B and F, and three independent experiments in panel A and D. ***p≤0.001, **p≤0.01, *p≤0.05, ns implies non-significant.

Since Notch1 could not protect Tregs in the absence of methionine, and SLC43A2 mRNA levels appeared to be Notch1 dependent (Fig. 3D), we asked whether Notch1 regulates SLC43A2 protein levels in Tregs, to thereby enable methionine-dependent survival. The levels of SLC43A2 and SLC7A5 protein in activated Notch1^+/+^ (Cre-ve) and Notch1^−/−^ (Cre+ve) Tregs were determined by Western Blot. Notch1^−/−^ Tregs showed a significant reduction in SLC43A2 protein levels relative to Notch1^+/+^ Tregs (Fig 4C(i)). However, no change was observed in SLC7A5 protein levels (Fig 4C(ii)). Further, freshly isolated Tregs showed decreased SLC43A2 protein levels in Notch1^−/−^ Tregs as compared to Notch1^+/+^ Tregs (Fig 4D) confirming that the difference in the SLC43A2 protein level is not due to differential activation in Notch1^+/+^ (Cre-ve) and Notch1^−/−^ (Cre+ve) Tregs. Since the non-nuclear Notch intracellular domain (NIC) protects Tregs from apoptosis in IL-2 limiting conditions (37), we examined the effect of NIC-NES on SLC43A2 protein levels. NIC-NES is recombinant NIC tagged to a Nuclear Export Signal to limit localization of NIC in the cytoplasm of Tregs (37). Overexpression of NIC-NES but not pBABE (control) by retroviral transduction restored the levels of SLC43A2 in Notch1^−/−^ Tregs (Fig 4E). Finally, we asked if the overexpression of SLC43A2 in Notch1^−/−^ Tregs could protect these cells from apoptosis triggered by IL-2 withdrawal. For this, activated Notch1^−/−^ Tregs were overexpressed with SLC43A2 and NIC-NES and pBABE vector control (Fig 4F(i)). Overexpressing SLC43A2 partially rescued Notch1^−/−^ Tregs from apoptosis (Fig 4F(ii)). The rescue by SLC43A2 was not equivalent to that provided by overexpression of NIC-NES (Fig 4F(ii)), suggesting additional factors in the survival program that are under Notch1 regulation.

Collectively, these data show that SLC43A2 is required for the methionine-dependent survival of Tregs in IL-2 deficient conditions, in a Notch1 dependent manner.

## Discussion

Understanding processes that enable death-survival decisions in T cells is a prerequisite to decipher how T cells regulate adaptive immune responses. Based on our data, we present a simple model for Tregs survival after IL-2 withdrawal (Fig 4G). Upon IL-2 withdrawal, Tregs require sustained uptake and transport of methionine, which is metabolized to SAM and utilized. To enable this, Notch1 (non-nuclear) functions to regulate the expression and activity of a specific solute carrier transporter, SLC43A2. SLC43A2 allows Tregs to take up methionine, and this can be subsequently utilized (Fig 4G). A reduction in SLC43A2 protein due to reduced Notch1 activity in these contexts abolishes Treg survival conferred by methionine. At this stage, these data suggest a coordinated ‘top-down’ (40) metabolic survival signaling and metabolic cascade in Tregs, where the IL-2 withdrawal and the activity of Notch1 together coordinate a methionine-dependent survival program.

In order to understand the metabolic programs that regulate cell death/survival/growth programs, it is useful to separate question into what metabolites are contextually needed and how these metabolic needs are sustained, versus what are the roles of specific metabolic programs in cells. Here, we identify methionine uptake and utilization as a limiting step in Tregs survival upon IL-2 withdrawal, with the SLC43A2 transporter sustaining this supply of sufficient methionine. The SLC class of transporters constitutes a large superfamily with increasingly important metabolic roles (41, 42), of which the more selective amino acid transporters (such as SLC7 or SLC43A) remain poorly studied. Currently, critical roles of SLC class transporters for other metabolites, particularly glucose import via GLUT or SLC2A transporters in controlling T cell metabolism to enable differentiation or activation are better known (43–46). As gatekeepers of nutrient (including amino acid) availability and therefore enablers of metabolic programs, it is likely that the amino acid transporting SLC proteins will also play critical roles in the survival or development of variety of cells. Understanding how these transporters function in different contexts, and how they are regulated by cytokine-dependent and nutrient signaling systems will be necessary to understand how unique metabolic programs are enabled in T cells. It will be just as important to understand how T cell fate regulating signaling systems such as mTORC1, Akt or Notch control the activity or functions of these transporters, to sustain specific metabolic programs that determine activation, differentiation or survival/death decisions in lymphocytes.

Finally, the specific functions of methionine in enabling Tregs survival remain unknown. Methionine, primarily through its metabolic product SAM, controls a variety of fundamental metabolic, signaling and regulatory processes (47–49). In a relevant example of T effectors, antigenic stimulation induced methionine uptake and utilization to drive a program of nucleotide and protein methylation for multiple fundamental processes (25), in contexts where growth and proliferation predominate (24, 50). Here, in a distinct context of Tregs survival (where there is no growth or differentiation), cells seem to similarly require substantial methionine uptake and consumption, to presumably enable a distinct survival program. Understanding these metabolic programs in Tregs for adaptive immune responses will be an exciting direction of future enquiry.

## Materials and methods

### Mice

C57BL/6J mice were procured from The Jackson Laboratory. The Notch1^lox/lox^ and Cd4-Cre∷Notch1^lox/lox^ (Notch1^−/−^) mouse strains were a gift from F. Radtke [École Polytechnique Federale de Lausanne (EPFL), Switzerland] (51). Animal experiments were approved by the Institutional Animal Ethics Committees of the National Centre for Biological Sciences (NCBS-TIFR) and inStem, Bangalore. The experiments were performed with mice of 6-12 weeks of age maintained per the norms of the Committee for the Purpose of Control and Supervision of Experiments on Animals, Government of India, as mentioned earlier (38). Colonies were tested for the full pathogen panel advised by the Federation of Laboratory Animal Science Associations.

### Cell lines

The HEK293T (HEK) cell line was obtained from American Type Culture Collection (ATCC) (Manassas, VA, USA). HEK and T cells were cultured and maintained as mentioned earlier (38). T cells were cultured in 5% heat-inactivated fetal bovine dialyzed serum (CMDS) whenever required for experiments. When required, T cells were cultured in RPMI without L-cysteine, L-cystine, L-Methionine (MP Biomedicals) and 5% heat-inactivated fetal bovine dialyzed serum (DOSAA).

### Reagents and antibodies

Reagents: Trizol (15596026) and SYBR™ Green Master Mix (Thermo Scientific, MA, USA), PrimeScript 1st strand cDNA Synthesis Kit (6110A) (Takara Bio, Shiga, Japan), Recombinant IL-2 (R&D Systems MN, Canada), Magnetic beads coated with anti-CD3 and anti-CD28 antibodies (Invitrogen, MA, USA), γ-Secretase Inhibitor X (GSI-X, 565771) and puromycin (508838) (Calbiochem-Merck Millipore Darmstadt, Germany), X-treme-GENE (6366236001), histopaque (10831) L-Ethionine (E1260), L-Cysteine (168149), and L-Methionine (M9625) (Sigma-Aldrich, St. Louis, MO, USA), 13C515N labeled L-methionine (CNLM-759-H-0.1) (Cambridge Isotope Laboratories, Inc., MA, USA). All other reagents were purchased from Himedia (India) or Sigma-Aldrich. L-Ethionine, L-Cysteine, and L-Methionine stocks were made in autoclaved distilled water, filter sterilized and stored at −80°C. Fresh aliquots were used for each experiment. Antibodies: PA5-71365 against SLC7A5 (Life technologies/Thermo Scientific); ab186444 against SLC43A2 and mN1A (128076) against Notch1 (Abcam, Cambridge, UK); ACTN05, MS-1295-P to Actin and MS-581-P0 to Tubulin (Neomarker, Fremont, CA, USA); Horseradish-peroxidase linked anti-mouse (7076S) and anti-rabbit IgG (7074P2) (Cell Signaling Technology, MA, USA).

### Plasmids

From Origene (MD, USA): Mouse shRNA plasmids to SLC43A2 (TR5143399) and SLC7A5 (TG513945), SLC43A2 Human Tagged ORF Clone (RC206639), and scrambled control. Plasmids pBABE and pBABE-NIC-NES were gifts from B.A. Osborne (University of Massachusetts/Amherst, MA, USA). SLC43A2 was subcloned into pBABE vector. Primers used for subcloning are listed in Table S2.

### Retroviral transductions

For retrovirus preparation, HEKs (0.25 × 10^6^) were seeded in 35 mm dishes (Greiner Bio-one, Kremsmünster, Austria). Cells were transfected the next day with plasmids containing the gene of interest (1.5 μg) and packaging vector pCL-Eco (1.5 μg for plasmids and 2 μg for shRNA) using X-tremeGENE. Retrovirus transduction was as described earlier (38). After transduction live cells were selected in histopaque (1.083 g/ml density) by centrifugation at 1500 rpm for 20 min. Cells were washed twice in RPMI-CM and PBS and cultured in media supplemented with IL-2 (1 μg/ml) and IL-7 (2 ng/ml) for another 22-24 h and used for apoptosis assays. Knockdown or overexpression of targeted genes was assessed by Western Blotting.

### Western Blot

Freshly isolated or activated Tregs (0.4-0.6 × 10^6^) were lysed and cell lysates were resolved by 10% SDS-polyacrylamide gel electrophoresis followed by western blot as described earlier (38). Primary antibody was used at the following dilutions: Notch1 (1:500), SLC43A2 (1:500), SLC7A5 (1:500), Actin (1:1000), and Tubulin (1:1000) diluted in 5% skimmed milk in TBST. Horseradish peroxidase-conjugated secondary antibody was used at 1:1000 dilution. The blots were developed using Super Signal West Dura substrate (Thermo Scientific), and images were acquired in iBright FL1000 Invitrogen.

### Isolation of Tregs

One and a half murine spleens were used to generate ~2×10^6^ natural Tregs (CD4^+^CD25^+^). Spleens were processed to obtain RBC-free lymphocytes as described earlier (38). Further, CD4^+^CD25^+^ natural Tregs were isolated from the splenocytes using the DynabeadsFlowComp Mouse CD4^+^CD25^+^ Treg cells Kit (11463D, Invitrogen) following the manufacturer’s instructions and as previously described (37)(38). Tregs were activated at a density of ~2×10^6^/ml with 20 μl/ml magnetic beads coated with antibodies to CD3 and CD28 in a 24 well plate. After 38-44 h of activation, Tregs were harvested from beads by magnetic separation, and the cells were further used for experiments.

### Metabolic profiling of Tregs

Tregs (2 × 10^6^, activated for 38 h) were cultured in CMDS without IL-2 for 1, 3, and 6 h. After the incubation, the cells were pelleted down at 2500 rpm for 3 min, and metabolites were extracted. Briefly, 1 ml of ice-cold 10% methanol was added without disturbing the pellet and further centrifuged at 2500 rpm for 3 min at 4°C. Further, 1 ml of 80% methanol (maintained at −45°C) was added to the pellet, vortexed for 15 s, and incubated at −45°C for 15 min. After the incubation, the tubes were vortexed for 15 s and centrifuged at 15000 rpm for 10 min at −5°C. The supernatant was transferred into fresh tubes without disturbing the pellet and centrifuged again at 15000 rpm for 10 min at −5°C. The supernatant was transferred to fresh tubes and dried using speedvac. The samples were stored at −80°C briefly, before analysis by targeted LC/MS/MS to assess methionine and other related metabolite amounts, using methods described earlier (52). To determine the uptake of methionine, Tregs (2 × 10^6^, activated for 38 h) were harvested and washed with PBS and further cultured in DOSAA and unlabeled methionine (150 μM) in the absence of IL-2 for 1, 3, and 6 h. 20 min before harvesting and metabolite extraction, the cells were incubated with 13C515N labeled methionine (150 μM). Subsequently, the relative amounts of labeled methionine, SAM or SAH were measured using targeted LC/MS/MS based approaches. MS-Q1/Q3 (Parent/Product) parameters used: methionine (Q1/Q3 150.1/56), SAM (Q1/Q3 399/250), SAH (Q1/Q3 385/136), 13C515N Methionine (Q1/Q3 155.1/60). Mass spectrometer used: AB Sciex 6500 QTRAP. Data normalization: Relative metabolite amounts are typically shown. For this, the first time point (T0) data is normalized to 1, and subsequent samples were compared relative to T0. Raw and normalized data are shown in Supporting worksheet 1.

### RT-PCR analysis

Tregs (4 × 10^6^, activated for 38 h) were harvested and cultured for 3 h without IL-2 in the presence or absence of GSI. After the incubation, cells were processed for RT-PCR using Maxima™ SYBR Green qPCR Master Mix and Bio-Rad CFX96 Touch™ Real-Time PCR Detection System as mentioned earlier (38). Relative change in transcript levels was calculated using the 2^−ΔΔCt^ method. Hypoxanthine-guanine phosphoribosyltransferase (HPRT) was used as the reference gene. The primers used for RT-PCR against murine genes are listed in Table S3.

### Apoptosis Assay

Tregs were washed with PBS thrice and 0.3 × 10^6^ cells/ml were cultured in a 48 well plate in the absence of IL-2 and with the required treatment for 18-22 h and processed for scoring apoptotic nuclei as described earlier (38). Approximately 200 cells were scored across several fields using fluorescent microscope (Olympus BX-60). To compare nuclear damage in Notch1^+/+^ (Cre-ve) and Notch1^−/−^ (Cre+ve; Cd4-Cre∷Notch1^lox/lox^) Tregs in the presence and absence of methionine, 1.2-1.4 x 106 were activated in a 48 well plate by co-culturing with beads coated with anti-CD3 and anti-CD28 antibodies (15 μl/ml). After 40-42 h, cells were separated from beads by a magnet and continued in culture in IL-2 (1 μg/ml) and conditioned media for another 20-24 h in a 24 well plate. Cells were further processed for apoptosis assay as described earlier (38).

### Statistical analysis and data presentation

Data are represented as Mean ± SD of two or three independent experiments (as indicated). Statistical significance was calculated using two-tailed Student’s t-test. Western blots were analyzed using ImageJ software and processed with Adobe Photoshop. Graphs and heat map were prepared using GraphPad Prism and figures were prepared using Illustrator.

## Acknowledgements

We acknowledge Freddy Radtke, EPFL, Switzerland for the Notch1^lox/lox^ and Cd4-Cre∷Notch1^lox/lox^ mice, Barbara A Osborne, Amherst USA for pBABE, and pBABE-NIC-NES plasmids, and I. Verma (Addgene, plasmid #12371) for the pCL-Eco construct. We acknowledge Bangalore Life Science Cluster (BLiSC), the Mass Spectrometry facility and Animal Care and Resource Center (ACRC) of NCBS-TIFR and DBT-inStem, Bangalore, India. Funding support: Department of Biotechnology, Government of India Grant BT/PR13446/COE/34/30/2015.

## Author Contributions

Designed experiments – AN, SP, PG, NS, AW, AS, SL; Performed experiments – AN, SP, PG, NS, AW, AD, UMV; analyzed and interpreted data – AN, SP, PG, NS, AW, AD, UMV, AS, SL; wrote the manuscript – AN, NS, AS, SL.

## Conflict Of Interest

The authors declare that they have no conflicts of interest with the contents of this article.

This article contains supporting information.

