## Supporting Information for "Methionine uptake via SLC43A2 transporter is essential for regulatory T lymphocyte survival"

**Materials included-** Table S1, S2 and S3, and excel worksheet 1.

**Table S1.**

| **Samples** |  | **Average Ct values** |  |
| --- | --- | --- | --- |
|  | **T0** | **No IL-2** | **No IL-2 + GSI** |
| HPRT | 23.76 | 24.40 | 24.16 |
| SLC3A2 | 21.39 | 21.81 | 21.41 |
| SLC43A1 | 24.57 | 24.80 | 25.38 |
| SLC43A2 | 22.46 | 22.17 | 23.05 |
| SLC6A17 | 33.34 | 33.30 | 32.14 |
| SLC7A5 | 21.10 | 21.45 | 21.32 |
| Notch1 | 21.34 | 21.99 | 22.22 |
| SLC1A5 | 32.82 | 33.59 | 33.65 |
| SLC7A8 | 29.34 | 30.05 | 29.48 |

**Table S1. Average Ct values used for Heat map.** Average Ct values of HPRT (reference gene used), indicated SLC transporters and Notch1 in Tregs at T0 and when cultured without IL-2 for 3 hours in the absence and presence (10 µM) of GSI. The Ct values were used to generate the Heat map in figure 3D.

**Table S2**

| **Restriction enzyme** | **Sequence** |
| --- | --- |
| SLC43A2 BamH1 | Forward:5'-CGCGGATCCATGGCGCCCACC -3' |
| SLC43A2 EcoR1 | Reverse:5'- CCGGAATTCTTACACGAAGGCCTCCT -3' |

**Table S2. Primers used for subcloning SLC43A2 into pBABE vector.**

**Table S3**

| **Gene** | **Forward (5’–3’)** | **Reverse (5’–3’)** |
| --- | --- | --- |
| HPRT | TCAGTCAACGGGGGACATAAA | GGGGCTGTACTGCTTAACCAG |
| Notch1 | ACAGTGCAACCCCCTGTATG | TCTAGGCCATCCCACTCACA |
| SLC3A2 | TGCTCAGGCTGACATTGTAGC | TCAGCCAAGTACAAGGGTGC |
| SLC43A1 | TTCACATGGTCTGGCCTGG | TGTGGTCCAAGGCTAACCC |
| SLC43A2 | ACAGTTTGGTAGCCTCACTGG | CCGGTAGCAGATGAGGTAAAGG |
| SLC6A17 | CCTTCATCAACTTCTTCACCTC | CGACCACACACTTCTCATTC |
| SLC7A5 | CTGGATCGAGCTGCTCATC | GTTCACAGCTGTGAGGAGC |
| SLC1A5 | TACATTCTGTGCTGCCTGCT | ATGAAACGGCTGATGTGC |
| SLC7A8 | CTAGCCTCCAATGCAGTTGC | GGCTCCAGCAAAGAACAGC |

**Table S3. Primers used for RT PCR against murine genes.**
